# EasyFlow: An open source, user friendly cytometry analyzer with graphic user interface (GUI)

**DOI:** 10.1101/2023.08.07.552387

**Authors:** Yitong Ma, Yaron Antebi

## Abstract

Flow cytometry enables quantitative measurements of fluorescence in single cells. The technique was widely used for immunology to identify populations with different surface protein markers. More recently, the usage of flow cytometry has been extended to additional readouts, including intracellular proteins and fluorescent protein transgenes, and is widely utilized to study development, systems biology, microbiology, and many other fields. A common file format (FCS format, defined by International Society for Advancement of Cytometry (ISAC)) has been universally adopted, facilitating data exchange between different machines. A diverse spectrum of software packages have been developed for analysis of flow cytometry data. However, those are either 1) costly proprietary softwares, 2) open source packages with prerequisite installation of R or Python and sometimes require users to have experience in coding or 3) online tools that are limiting for analysis of large data sets. Here we present EasyFlow, an open source flow cytometry analysis GUI based on Matlab or Python, that can be installed and run locally cross-platform-ly (Windows and MacOS), without prerequisite user having previous knowledge on coding. The python version (EasyFlowQ) is also developed on a popular plotting framework (Matplotlib) and modern user interface (UI) toolkit (Qt), allowing more advanced users to customize and keep contributing to the software, as well as its tutorials. Overall, EasyFlow serves as a simple-to-use tool for inexperienced users with little coding experience to use locally, as well as a platform for advanced users to further customize for their own needs.

## Background

Flow cytometry enables quantitative high-throughput measurements of fluorescence in small particles and single cells, and has been widely used in immunology, microbiology, synthetic biology and many other fields. Different flow cytometry machines were developed over the years, with distinct features and utilizing different technologies [1]. However, the FCS file standard (defined by the International Society for Advancement of Cytometry (ISAC) [2,3]) has been universally adopted as a common output format across all devices. This standardization enormously facilitates the data exchange between different machines and labs.

A diverse spectrum of software solutions has been developed to analyze flow cytometry from FCS files. When comparing these tools, several key features are important to highlight. First, the analysis of flow cytometry data depends heavily on visual user interactions. This includes viewing different histograms or scatter plots and determining visually gating strategies to extract specific subpopulations of cells. In addition, while a core set of operations are used routinely, flexibility in the analysis and user customization are important as well. Finally, platform independence, availability, maintenance, and cost of the software are additional aspects that differ between existing solutions. Current tools can be roughly categorized into three categories - proprietary software, open-source libraries, and online analysis tools. Proprietary software, e.g., FlowJo (BD Bioscience) and FCS Express (De Novo Software), provides full-feature powerful analyzing tools. However, such software can be costly and lack transparency and flexibility for user customization. Notably, there are several tools with graphical user interfaces (GUIs), including Flowing Software, Cyflogic, and FCSalyzer, that are freely available for academic users. However, none of them is open-source or capable of further modification and transparency. Some of them are not actively maintained, and they are often operating only on specific platforms (e.g., Windows only). Open-source libraries (usually written in R or Python) provide superior flexibility but require users to have previous, and usually significant, experience in coding. A major weakness of these tools is the lack of visual interactions that are required for many common routine analysis tasks. Online analysis tools can be either commercial (e.g., CellEngine by Cell Carta) or freely available (e.g., Floreada.io). These tools are easy to use across platforms and require no installation at all. However, big datasets and limited internet connections present significant challenges to the routine use of these tools.

Here we present the matlab based EasyFlow (github.com/AntebiLab/easyflow) and its derivative standalone Python EasyFlowQ (ym3141.github.io/EasyFlowQ/), which are open source, utilize user-friendly GUI, can be run on multiple platforms (Windows, MacOS and Linux), and requires no coding knowledge. Both versions were designed to enable a simple tool for viewing, gating and analyzing flow data, while allowing further higher-level analysis using the full power of the corresponding scripting language. Being open source, these tools provide transparency of data processing and allows further customization and collaborative improvements for advanced users.

## Results

EasyFlow is an open source software designed as an intuitive tool for visual based analysis of single cell flow cytometry data. Its main goal is to simplify the identification and extraction of the proper subset of events for further analysis, as well as generate the standard histogram and scatter plots. Importantly, it provides the capacity to extract the resulting data, and key statistical parameters for downstream secondary analysis. To serve this goal, EasyFlow utilizes a simple, plot-centric user interface (Figure 1A and 1B) featuring a single plotting region at the center of the GUI, with most used functions, including sample management, channel selection, gating, plotting options etc. exposed on the top layer. EasyFlow supports the basic functions for analyzing FCS files: loading and parsing FCS files; plotting histogram, scatter, density and contour (matlab version only currently); visually creating and editing 2D and 1D gates; exporting raw data, auto and manual compensation, statistical analysis, etc. We also programmed convenient features such as batch sample renaming and session saving. Statistical features for each gated population can be computed and exported to further analysis. EasyFlow is implemented in both matlab and python programming environments, providing three main characteristics: capacity for exporting and interaction with raw data, cross-platform and standalone capability, and flexibility for further community development.

**Figure 1:**
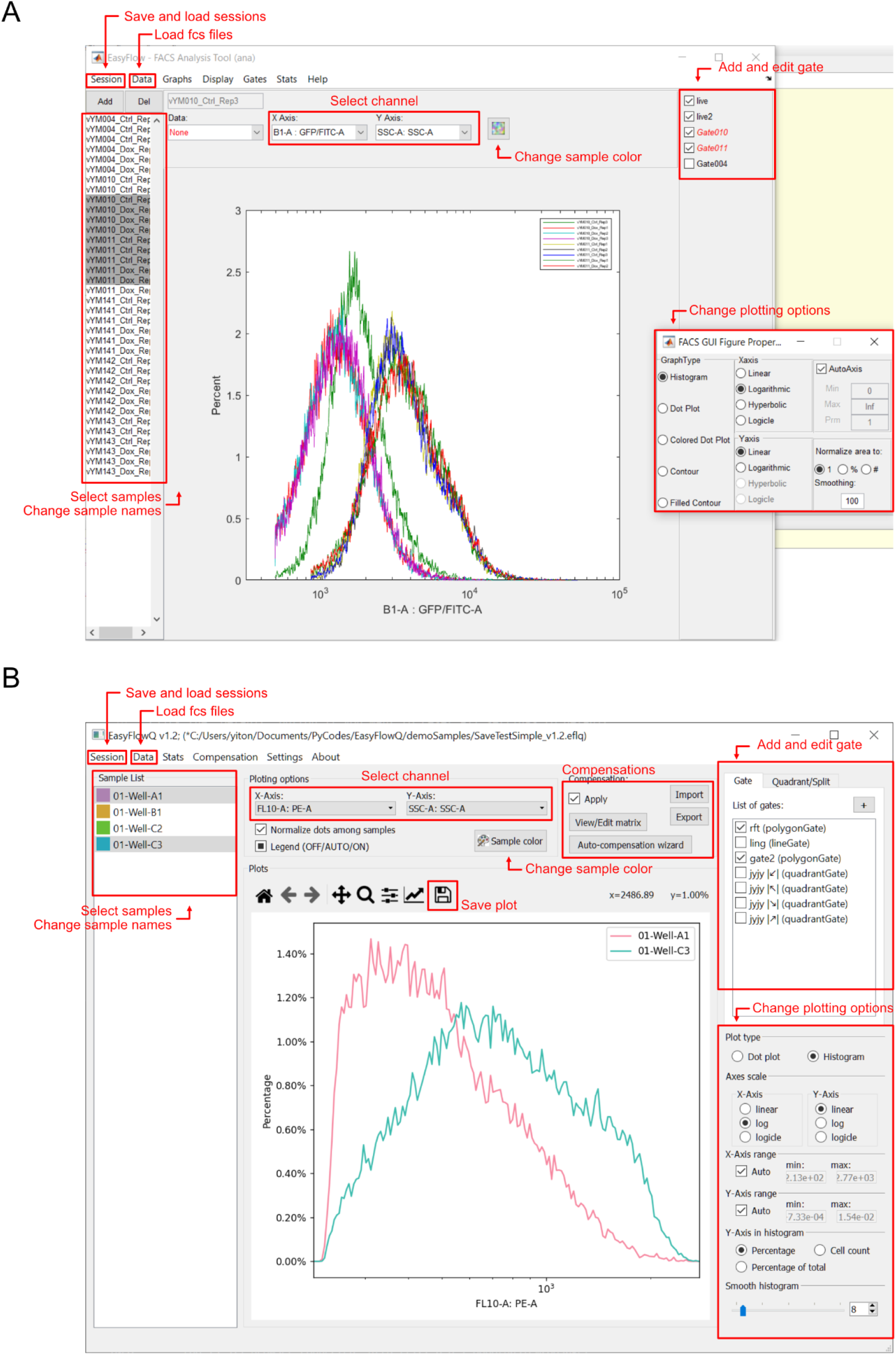
Screenshots of the EasyFlow (Matlab) (A) and EasyFlowQ (B), with basic functions including managing FCS files, plotting and gating, annotated.

### Exporting and interaction with raw data

In addition to exporting statistics, we designed both versions of EasyFlow to be able to export the processed data (e.g. gated samples) for further in-depth processing. In EasyFlow(Matlab), data can be exported into the Matlab working space for further processing with Matlab scripts. In EasyFlowQ, the data are exported as comma-separated values (.csv) files, that can be conveniently accessed by commonly available programs (e.g. Excel) or programming scripts (Python and R). These abilities allow users to further analyze their data with great flexibility.

### Cross-platform and standalone capability

EasyFlow was designed for cross-platform usage. The original EasyFlow (Matlab) runs on standard Matlab installation (versions later than 2017a). EasyFlowQ (Python) runs on standard Anaconda installation (based on Python 3, version later than 3.7 recommended). Both software are available on all three major operating systems (Windows, MacOS, and Linux). Additionally, we packaged EasyFlowQ using Pyinstaller and InstallForge (Windows) or create-dmg (MacOS) into installers, to further eliminate the requirement of installing additional software (packages). These installers can be used on either Windows or MacOS to create a standalone software, with a similar experience as most mainstream applications.

### Flexibility for further community development

We built both versions of EasyFlow to allow further development from community contribution. More specifically, we ensured that the source code can be run by standard installation of Matlab (EasyFlow) or the popular python installation Anaconda in its default state (EasyFlowQ). For the latter, we further provided a list (yaml file) of recommended packages that can be “imported” by existing Anaconda installation. We also used Matplotlib and PyQt5, two of the most popular packages in the Python programming community, as plotting and UI framework respectively. Furthermore, both versions of EasyFlow are hosted on Github, making them possible to benefit from the well-established, community driven, open source software developing model.

## Conclusion

EasyFlow provides an accessible and simple solution for analysis of flow cytometry data. It enables standard analysis tools and provides the capacity for in depth analysis using scripting languages such as Matlab or Python. For everyday users looking for an easy-to-use local FCS analyzer, we recommend EasyFlowQ (python based), as its standalone packages require no pre-installation of either Matlab or Python, as well as have a user experience more similar to other native Windows or MacOS applications. EasyFlow is optimized for users that are familiar with Matlab, and need access to the raw data for subsequent analysis with Matlab script. Contributions through Github, including forking, reporting bugs and feature requests are welcome.

## Acknowledgment

We would like to thank Nir Friedman, Inbal Eizenberg and Eric Shifrut for comments and suggestions on early versions of EasyFlow; Michael Elowitz for funding and mentorship; Evan Mun and Kaiwen Luo for sharing their data for testing the program. This work is supported by the National Institutes of Health grant RO1 HD075605A and R01 MH116508, and by National Science Foundation grant EF-2021552 under subaward UWSC10142.

